# Low incidence of plastic ingestion among three fish species significant for human consumption on the island of Newfoundland, Canada

**DOI:** 10.1101/332858

**Authors:** Max Liboiron, Jessica Melvin, Natalie Richárd, Jacquelyn Saturno, Justine Ammendolia, France Liboiron, Louis Charron, Charles Mather

## Abstract

This study reports the first baselines of plastic ingestion for three fish species that are common food fish in Newfoundland, Canada. Species collections occurred between 2015-2016 for Atlantic cod (*Gadus morhua*), Atlantic salmon (*Salmo salar*), and capelin (*Mallotus villosus*). The frequency of occurrence (%FO) of plastic ingestion for both spawning Atlantic salmon (*n*=69) and capelin (*n*=350) was 0%. Of the 1,010 Atlantic cod collected over two years, 17 individuals had ingested plastics, a %FO of 1.68%. This is the only multi-year investigation of plastic ingestion in Atlantic cod for the Northwest Atlantic, and the first for capelin and salmon in the region. Considering the ecological, economic, and cultural importance of these fish species, this study is the beginning of a longitudinal study of plastic ingestion to detect future changes in contamination levels.

**Highlights:**

- Ingestion rate (%FO) of plastics in Atlantic cod is 1.68%
- Ingestion rate (%FO) of plastics in Atlantic salmon and capelin is 0%
- First study of plastic ingestion rates in Atlantic salmon and capelin
- Multi-year baseline of plastic ingestion in Atlantic cod in the Northwest Atlantic
- Plastic ingestion rates for three food fish species in Newfoundland, Canada, are low

**Terms:**

- Frequency of occurrence (%FO): the number of individuals in a population or group that have ingested plastics (not indicative of the number of particles ingested per individual)

## Introduction

Since its rise as a consumer and commercial material in the 1940s (Meikle, 1997), plastic production has increased exponentially, giving rise to an estimated production of 400 million tonnes in 2017 (Hopewell et al., 2009; Plastics Europe, 2016). This growing rate of plastic production has been reflected by increasing levels of plastic debris in the marine environment (Barnes et al., 2009; Cózar et al., 2014; Thompson et al., 2004). Surveys from remote areas report marine plastics in diverse ecosystems, including the Arctic, Antarctica, and isolated islands of the Southern Ocean and South Pacific (Eriksson and Burton, 2003; Lavers and Bond, 2017; Mallory, 2008; Zarfl and Matthies, 2010). Yet, remote, northern regions can still constitute a blind spot in marine plastics monitoring for a variety of reasons, including high costs, seasonal restrictions, a lack of research infrastructure, and a lack of standardized monitoring procedures adapted to harsh environments (Bergmann et al., 2017; McWilliams et al., 2017; Poon et al., 2017).

The northeastern-most Canadian province of Newfoundland and Labrador (NL) comprises of a section of coastal mainland (Labrador) and an island (Newfoundland). The province of NL is the most sparsely populated province in Canada, with a population density of 1.4 persons/km^2^ (Statistics Canada, 2012). Much of this human population is concentrated along the coastline with access to fisheries. To date, only a handful of studies have examined the marine plastic landscape surrounding Newfoundland, despite the province’s high degree of marine industrial activity and its dependence on the ocean for food, culture, and livelihoods (Canadian Association of Petroleum Producers, 2017, 2016; Fisheries and Oceans Statistical Services, 2016; The St. Lawrence Seaway Management Corporation, 2016). Generally, remote islands are characterized by a high representation of plastic debris from sea-based sources, most commonly through fishing and shipping activities (UNEP, 2009). These activities are expected to be significant contributors to the marine plastic landscape around the island of Newfoundland, considering the waters surrounding the island contribute over 80% of the national fisheries landings (Fisheries and Oceans Canada, 2017a) and facilitate the movement of over 160 million tonnes of shipping cargo annually (The St. Lawrence Seaway Management Corporation, 2016).

Despite the low human population, land-based sources of marine plastics should also be considered in Newfoundland. It is estimated that the province generates 493,595 metric tonnes of solid municipal waste per year (Auditor General of Newfoundland and Labrador, 2014). In 2016, a provincial highway litter audit reported that 42.7% of all collected litter from across the province were of plastic origin (13,745 of 32,190 pieces) (Multi-Materials Stewardship Board, 2016). Terrestrial waste can be transferred to the marine environment through numerous pathways typical of coastal regions such as wind and run-off, but some local waste management practices may increase these rates in NL. Most rural municipal sewerage is not treated in the province, and 760 sewer outfalls release 39,000 m^3^ of effluent daily (Government of Newfoundland and Labrador, 2016). Small-scale incineration and open burning of waste, as well as direct dumping remain an issue in rural, outport communities (Avery-Gomm et al., 2016; Government of Newfoundland and Labrador, 2017; Henemen, 1988).

### The significance plastic ingestion by fish in Newfoundland

Plastic ingestion by fish is a primary concern in Newfoundland. Since over 90% of surface water marine plastics are less than 5 millimeters in size (Eriksen et al., 2014), they have a high degree of bioavailability to a wide range of trophic levels in the marine food web. Marine plastics are associated with contaminants that can take the form of ingredients and by-products of the plastic material itself (such as UV stabilizers, softeners, flame retardants, non-stick compounds and colourants), as well as contaminants adsorbed from the surrounding seawater (such as PCBs and DDT) (Lithner et al., 2012; Mato et al., 2001; Rochman et al., 2015). The accumulation of toxic chemicals in marine species can be transferred up the food chain via biomagnification, thus potentially negatively affecting apex predators like humans (Ochi, 2009; Teuten et al., 2009; Browne et al., 2013; Rochman et al., 2013; Koelmans et al., 2016). Industrial chemicals associated with marine plastics have been correlated to negative health effects in humans, including endocrine disruption, heart disease, and developmental disorders (Colborn et al., 1994; Kim et al., 2002; Lang et al., 2008; Melzer et al., 2010). Despite these preliminary links further research is required to better understand the full effects that plastics consumed at lower trophic levels in the food web can have on top consumers (Cole et al., 2011; Engler, 2012). Nonetheless, the burden of polluted wild food sources disproportionately affects rural and low-income communities where country foods are relied on for sustenance (Shepard et al., 2002).

Despite the devastating collapse of the Atlantic cod (*Gadus morhua*) stock in Newfoundland and the resulting moratorium on its fishery in 1992, fishing - both “recreational” (sustenance) and commercial - remains financially and culturally important to the people of Newfoundland. In 2016, commercial fisheries employed over 17,000 individuals and landed over 220,000 tonnes of catch (fish and shellfish) resulting in a production value of $1.4 billion (Fisheries and Land Resources, 2017). Within the context of the Newfoundland fishery, Atlantic cod yielded the second highest landed value for all captured groundfish species in the country (∽$23 million), while capelin (*Mallotus villosus*) yielded the highest landed value of all pelagic species (∽$13.5 million) (Fisheries and Oceans Canada, 2017b). The provincial recreational fisheries have one of the highest participation rates in Canada with 71,382 participants in 2010 (Fisheries and Oceans Canada, 2012a). Most of this participation comes from the recreational groundfish fishery for Atlantic cod that does not allow the selling of catch (Fisheries and Oceans Canada, 2012a; Government of Canada, 2017). Fishers retain their catch (as opposed to catch and release), and fish are relied on for sustenance (Fisheries and Oceans Canada, 2012a). In a study of seafood preferences in rural communities in Western Newfoundland, both Atlantic cod and Atlantic salmon were the species most commonly consumed by respondents (81% and 42%, respectively), while capelin was eaten less often, but remained a part of the diet of a majority of respondents (Lowitt, 2013).

This research provides a baseline for plastic ingestion in these three common Newfoundland food fish; Atlantic cod, Atlantic salmon, and capelin. This research: (1) builds upon and validates the existing preliminary finding of a %FO of 2.45% in Atlantic cod collected in 2015 from the eastern coast of Newfoundland (Liboiron et al., 2016) to generate a validated, multi-year baseline for plastic ingestion in Atlantic cod of the Northwest Atlantic; and (2) establishes baselines of plastic ingestion in Atlantic salmon and capelin on the island of Newfoundland. Based on a review of English-language, published, scientific literature, no previous studies on the ingestion of plastics by Atlantic salmon or capelin exist. Other studies of plastic ingestion in Atlantic cod in the Northeast Atlantic have been conducted the North and Baltic seas (Bråte et al., 2016; Foekema et al., 2013; Lenz et al., 2015a; Rummel et al., 2016). The production of a baseline of plastic ingestion levels in these three species is the first step towards a reliable and useful long-term monitoring program for the region, especially given their cultural, ecological, and economic value on the island of Newfoundland.

## Methods

### Collection protocols

This study synthesises plastic ingestion data from different projects conducted around the island of Newfoundland between 2015 and 2016. Accordingly, the geographical range of sampling covered much of the island’s inshore bays and harbours, as well as offshore regions in the Gulf of St. Lawrence, the Grand Banks, as well as a collection site north of Labrador (Davis Strait) (Fig. 1). “Offshore” was considered as any region >30 km from the shoreline whereas “inshore” was an area <30 km from the shoreline (30 km was selected based on the distance indicated by fishers whose activities are governed by these categories). Recreational fishers generally captured fish within 10 km of the shoreline, while commercial and scientific collections almost exclusively captured fish from at least 30 km offshore. Fish were collected using one of two methods; (1) collaborations with existing scientific surveys (SS) offshore (Atlantic cod, Atlantic salmon and capelin), and (2) citizen science surveys (CSS) of inshore recreational fisheries (Atlantic cod). Gastrointestinal (GI) tracts were removed in the field for both Atlantic cod and Atlantic salmon, while capelin were collected whole. Collection methods within the study are outlined in Table 1. For further details regarding collection protocols refer to S1.

**Table 1.** Summary of collection details for the Atlantic cod, Atlantic salmon and capelin examined for plastic ingestion rates.

| Collection details | Atlantic cod<br>(*Gadus morhua*) |  |  | Atlantic salmon<br>(*Salmo salar*) | Capelin<br>(*Mallotus villosus*) |
| --- | --- | --- | --- | --- | --- |
| **Sample size (n)** | 204 | 348 | 458 | 69 | 350 |
| **Collection date (month, year)** | October 2015 | July - October 2016 | April - June 2016 | February - October 2016 | July 2015 |
| **Collection method** | Citizen Science Survey | Citizen Science Survey | Scientific Survey | Citizen Science Survey | Scientific Survey |
| **Environment** | Marine | Marine | Marine | Freshwater | Marine |
| **Sample sites** | (1) Quidi Vidi<br>(2) Petty Harbour<br>(3) Portugal Cove - St. Philip's | (1) Quidi Vidi<br>(2) Petty Harbour<br>(3) Portugal Cove - St. Philip's<br>(4) Witless Bay<br>(5) Bauline East<br>(6) Brigus South | NAFO divisions: 3Ps, 3O, 3N, 3L, 0B<br>(1) Placentia Bay<br>(2) St. Pierre Bank<br>(3) Fortune Bay<br>(4) Burgeo Bank<br>(5) Green Bank<br>(6) Halibut Channel<br>(7) Southern Grand Banks<br>(8) Eastern Grand Banks<br>(9) Davis Strait | Campbellton River | NAFO divisions: 3Ps, 3L, 3K<br>(1) Bonavista Bay<br>(2) Conception Bay<br>(3) White Bay<br>(4) Notre Dame Bay<br>(5) Trinity Bay<br>(6) Lawn |
| **Variables recorded** | Food presence<sup>a</sup><br>Body length (n=38)<br>Sex (n=38)<br>Sample site<br>Inshore/ offshore<sup>c</sup><br>Date | Food presence<sup>a</sup><br>Body length (n=325)<br>Sex (n=287)<br>Sample site<br>Inshore/ offshore<sup>c</sup><br>Date | Food presence<sup>a</sup><br>Sex (n=445)<br>Depth of collection<sup>b</sup><br>Sample site<br>NAFO division<br>Inshore/ offshore<sup>c</sup><br>Date | Food Presence<sup>a</sup><br>Body length<br>Sex<br>Sample site<br>Date | Food Presence<sup>a</sup><br>Body length<br>Sample site<br>NAFO division<br>Date |
| **Plastic identification methods** | Visual | Visual<br>Raman Spectrometry | Visual<br>Raman Spectrometry | Visual<br>Raman Spectrometry | Visual<br>Raman Spectrometry |

<sup>a</sup>Food was considered present if prey items were found in the GI tract (stomach and/or intestines)
<sup>b</sup>Depth of collection (range: 46-442m) data points were only obtained for Atlantic cod collected via Scientific Survey (SS) (n=458)
<sup>c</sup>Offshore collections occurred in any region >30 km from the shoreline whereas inshore was an area <30 km from the shoreline

**Fig. 1.**
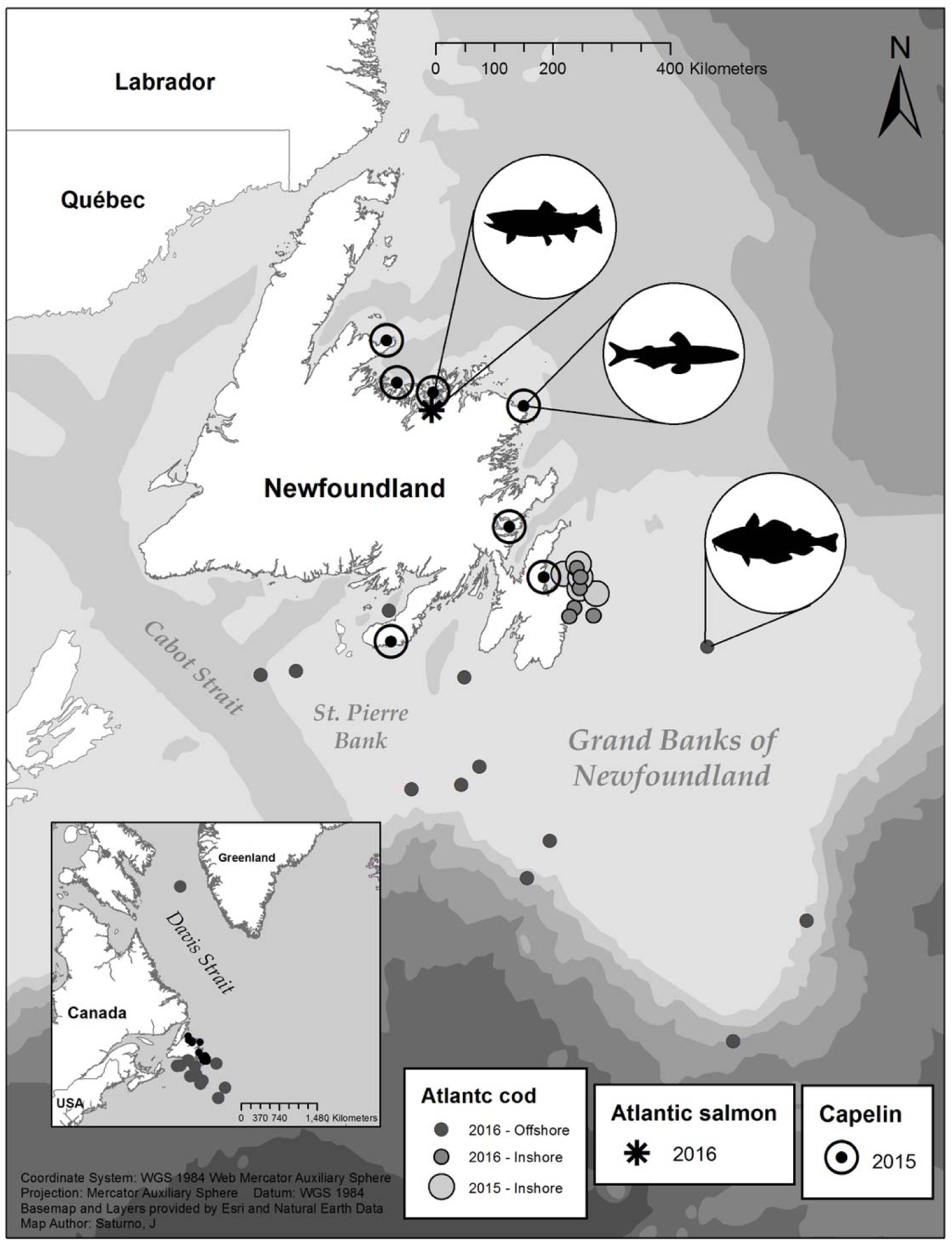
Collection locations for the Atlantic cod, Atlantic salmon, and capelin from 2015-16 (map: J. Saturno).

## Laboratory procedures

### Dissection of GI tracts

We used standardized laboratory plastic ingestion protocols identical to Liboiron et al., (2016) to compare previous studies in the province, which in turn have been modified from those used in bird ingestion studies by van Franeker et al., (2011). GI tracts (or whole fish in the case of capelin) were thawed for at least 2 hours prior to dissection. Once thawed, an incision was made along the entire length of the tract, from the esophagus to anus. A double sieve method was used for analyzing Atlantic cod and salmon by rinsing and scraping GI tract contents onto stacked sieves with mesh sizes of 4.75 mm (#4) and 1 mm (#18) to aid in sorting items. Due to the small size of capelin and their natural prey items (zooplankton), a single 1 mm sieve was used for capelin GI tracts. Once digestive contents were transferred to sieves, the GI tracts were visually and tactilely inspected for imbedded debris and thoroughly rinsed over the sieves. Sieve contents were continuously and gently rinsed with cold water so any organic and inorganic debris were removed from plastic particles to enable proper identification (Löder and Gerdts, 2015). Any particles suspected to be non-food items (anthropogenic or not) were removed and transferred to a petri dish for further analysis. The presence or absence of prey/digestive content in the stomach and/or intestines was noted. Although rare, there were some instances of relatively intact fish in the GI tract contents that were viable for secondary plastic ingestion analysis (*n*=5) but were too degraded for a positive species identification. These prey fish were analysed following the same methods described above in an attempt to identify the secondary ingestion of plastics.

### Contamination

Plastic contaminants were occasionally found on the external surface of the stomach and/or intestines for Atlantic cod (*n*=15). These particles were identified as wood splinters and paint chips that visually matched the inventory of field tools and were assumed to have been transferred during sample collection. To control for microplastic contamination in the laboratory, we used a cotton laboratory coat, wore gloves, tied hair back, rinsed all tools under tap water before and after each sample, and used daily control sticky dishes to capture particles settling from the air. In the event that a fibre was found in a GI tract, it was compared to fibres present in the control dish.

### Quantification of plastics

Visual identification of microplastics through use of a microscope has been confirmed as a reliable method for identifying particles greater than 1 mm in size (Song et al., 2015). As such, in deference to a protocol that could be used by citizen scientists, the lower limit of plastics identifiable by this research is restricted to 1 mm in the largest dimension.

Each particle that was identified as a probable plastic was individually wrapped in filter paper and dried for at least five days. Once dried, particles were visually examined under a stereomicroscope (Olympus SZ61 stereomicroscope, maximum magnification 45X), and a compound microscope (Ecoline, Eco Series, maximum magnification 100X). Particles identified as plastic at this stage were set aside to be verified by Raman micro-spectrometery (see section below).

Plastic ingestion is most commonly quantified in the literature as a frequency of occurrence (%FO), a metric that is indicative of the number of individuals in a sample population that ingested plastic(s) (Cannon et al., 2016; Peters et al., 2017; Provencher et al., 2016; Rochman et al., 2015; Romeo et al., 2015; Rummel et al., 2016). Following this metric, we recorded and quantified the presence or absence of plastic(s) for each individual fish and calculated the population average (%FO). The colour, weight, length, type and weathering details of plastics were recorded. Plastics were weighed on a scale sensitive to 0.0001 g and length measurements were taken from the longest axis of the plastic using digital calipers sensitive to 0.01 mm. There remains a debate in the literature over the designation of plastic size classes in terms of micro, meso, and macro designations (GESAMP, 2015). Following the designations used by Bråte et al. (2016) and Cannon et al. (2016), and the recommendation of Provencher et al. (2016) for standardization, plastics smaller than 5 mm in maximum length were classified as microplastics, while plastics from 5-20 mm were classified as mesoplastics (and those over 20 mm as macroplastics, though such plastics are rare in ingestion studies). When plastics were measured they were not stretched or manipulated to facilitate a longer axis, but were measured according to their size at time of recovery (Avery-Gomm et al., 2016; Liboiron et al., 2016). The size of ingested plastics is important to record because it is necessary to evaluate how much surface area plastics occupy in the GI tract and the surface area available for the transfer of contaminants (Jabeen et al., 2017; Mato et al., 2001; Provencher et al., 2016; Rummel et al., 2016; Silva-Cavalcanti et al., 2017).

Plastic types were categorized as either: industrial resin pellets, sheet/film (i.e. plastic bags), threads (i.e. fishing line, rope), foamed plastics (i.e. polystyrene packaging), fragments (i.e. hard plastics), fibres (i.e. from clothing), microbeads (i.e. from personal care products) or other. In order to evaluate the state of plastic erosion each particle was assessed for: discolouration, fraying, fracturing, pits and grooves, and adhered particles (following Corcoran et al., 2009), as well as burning or melting (following Avery-Gomm et al., 2016).

### Raman micro-spectrometry

Raman micro-spectrometry was conducted on particles obtained from: Atlantic cod collected in 2016, Atlantic salmon and capelin. This analysis was not available for the Atlantic cod collected in 2015. However, no plastics from 2015 were visually ambiguous. Particles were washed in ethanol and allowed to dry in preparation for Raman micro-spectrometry. Samples were then placed on a silica wafer of known Raman spectrum (520 cm^−1^ peak) and analysed using a Raman micro-spectrometer (Reinshaw InVia with 830 nm excitation) set at a 20x Olympus objective. To ensure samples were not burnt during the processing, laser power did not exceed 5%, and in cases of high fluorescence (in 4/5 samples), laser power was reduced to 1%. Results were analysed by using WiRE 3.4 software. A reference spectra was used to compare the Raman spectrum of each experimental sample to the following common marine plastic polymers: acrylonitrile butadiene styrene (ABS), cellulose acetate, polyamide (PA), polycarbonate (PC), polyethylene (PE), polyethylene terephthalate (PET), poly (methyl methacrylate) (PMMA), polypropylene (PP), polystyrene (PS), polyurethane (PU) and polyvinylchloride (PVC) (Bråte et al., 2016; Engler, 2012; Lenz et al., 2015b; Plastics Europe, 2016).

## Data Analysis

When frequency of occurrence for a species was >0% (Atlantic cod), the relationship between %FO and the following variables were evaluated: 1) food presence, 2) sex, 3) depth of collection, 4) sample site, 5) NAFO management area, 6) distance offshore, 7) day of year (DOY), 8) month, and 9) year. Each variable was modelled using a generalized linear model with a binomial error structure. Assumptions of residuals homogeneity, independence and normality were visually tested and met for all models using lag plots, residual versus fit plots, and quantile-quantile (QQ) plots. Models were then tested with a ANOVA for hypothesis testing; the null hypothesis was rejected when *p*<0.05 (95% confidence interval). All analyses were carried out using R Studio version 1.0.143. All graphs were generated using Prism Graphpad version 7.0 (Prism and Graphpad, 2016) and maps were generated using ArcGIS version 10.5.1 (ESRI, 2014).

Plastic characteristics (type, colour, length, mass, weathering and polymer) were quantified as a percentage of all particles. The Joint Research Centre of the European Commission recommends that for the standardization of plastic ingestion research, in addition to the %FO, the following metrics be reported: “1) abundance by number (average number of items per individual), and 2) abundance by mass (weight in grams, accurate to 4^th^ decimal)” (Galgani et al., 2013, p. 78). Both of these metrics were calculated for the entire sample population of Atlantic cod (*n*=1,010), as well as for the sub-population of individuals that had ingested plastic (*n*=17). Particles that could not be weighed (because they were lost prior to weighing or were too small to register) were assigned a mass of 0 g for calculations involving the entire sample population, and removed for calculations for the sub-population of individuals with ingested plastics.

## Results

A total of 1,429 individuals from 3 species were collected and plastics were found to have been ingested by 17 individuals (all Atlantic cod), resulting in a multi-species %FO of 1.19% (*n*=1,429, SD=0.110). Although %FO of plastics is commonly reported for multi-species populations (Anastasopoulou et al., 2013; Davison and Asch, 2011; Neves et al., 2015; Phillips and Bonner, 2015), this study will focus on %FO for separate species so that trends can be clearly examined (Table 2). This is particularly important in species with 0%FO, to avoid false connotations of ubiquitous plastic consumption by all fish species (Liboiron et al., 2018).

**Table 2.** Summary of the results for plastic ingestion rates (%FO) among three fish species collected from Newfoundland.

| Species | Collection year(s) | Sample size ($n$) | Mean plastic ingestion rate (%FO±SD) |
| --- | --- | --- | --- |
| Atlantic cod<br>(*Gadus morhua*) | 2015-2016 | 1,010 | 1.68 ± 0.129 |
| Atlantic salmon<br>(*Salmo salar*) | 2016 | 69 | 0 |
| Capelin<br>(*Mallotus villosus*) | 2015 | 350 | 0 |

### Atlantic cod

A total of 1,010 Atlantic cod were collected and 17 fish were found to have ingested plastic, a %FO of 1.68% (*n*=1,010, SD=0.130). None of the GI tracts found to contain plastic had any holes or perforations. One GI tract that contained plastic had green paint from a green splitting table on the exterior wall prior to dissection; this paint was discounted as contamination. All but two of the 17 fish with ingested plastic (*n*=15) had ingested only one plastic, while two of the fish collected in 2015 had ingested two plastics each. The mean number of plastics ingested per individual for the entire sample was 0.019 particles/individual (*n*=1,010, SD=0.150). Of the sub-population of Atlantic cod that had ingested plastic, the mean number of plastic particles ingested per individual is 1.12 particles/individual (*n*=17, SD=0.330).

The %FO of the 2015 citizen science survey collection did not significantly differ from individuals collected in 2016 (*p*=0.730, df=1, *n*=552). Therefore, the subsequent analyses of these datasets were combined (with the exception of the analysis of year as an independent variable) and are hereafter referred to as the citizen science survey (CSS). A total of 12 of 552 fish from the CSS of Atlantic cod had ingested plastic, yielding a %FO of 2.17% (*n*=552, SD=0.150). The scientific survey (SS) of Atlantic cod found plastic in 5 out of 458 fish collected for a %FO of 1.09% (*n*=458, SD=0.100). The comparison of the CSS and SS collections yielded no significant difference in plastic ingestion (*p*=0.170, df=1, *n*=1,010), and are therefore presented together as a total %FO (1.68%) for the species in Table 2.

The results of binomial generalized linear model fits of plastic ingestion are presented in Table 3. Notably, both food presence and sex were each significantly related to %FO (*p*<0.05), while there was no evidence of a significant relationship between %FO and temporal or regional variables (*p*>0.05). All 17 individual cod that had ingested plastic had also ingested other food items. Although most individuals had consumed food (71%), the presence of food in the stomach was a significant factor in positively predicting the presence of ingested plastic (*p*<0.01), whereby individuals that ingested prey were more likely to contain plastic. Sex was identified for 10 out of the 17 individuals with ingested plastic. Of these 10 individuals, 9 were females. Females were determined to have a significantly higher %FO than males (*p*<0.05).

**Table 3.**
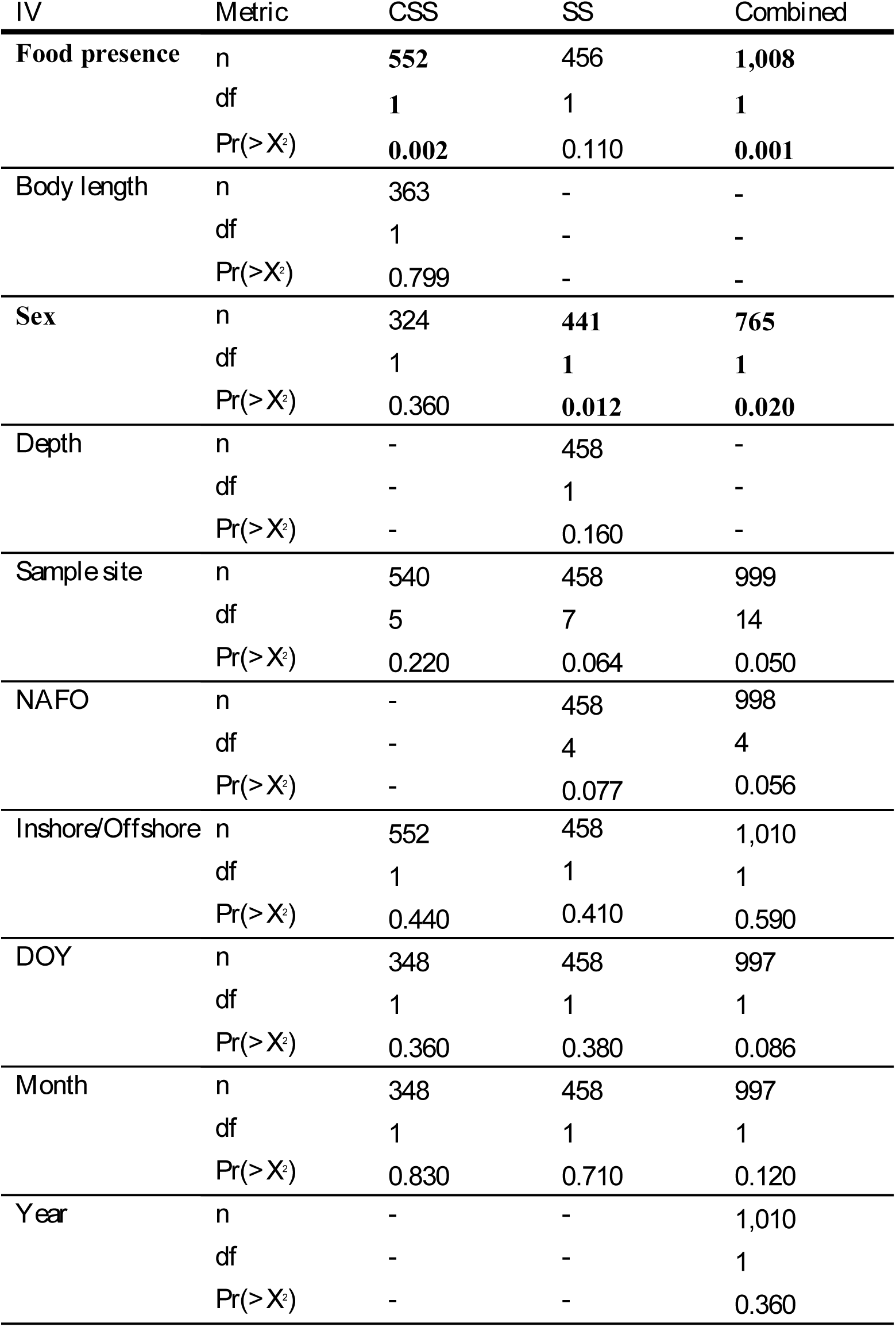
Independent variables found that had a statistically significant effect (indicated by bolding) on the %FO of plastic ingestion in Atlantic cod (Atlantic salmon and capelin were not tested, due to an absence of ingested plastics). Dashes represent the scenario in which there was not enough data present to conduct analysis. [IV=explanatory variable; CSS=2015-2016 citizen science surveys; SS=scientific surveys]

### Atlantic salmon

None of the 69 Atlantic salmon that were caught in 2016 had ingested any plastics, yielding a %FO of 0% for the species. Notably, only 20% of individuals had ingested food. This proportion was considerably less than what was found in Atlantic cod (see above), though salmon are caught while spawning, while cod are not.

### Capelin

Of the 350 capelin collected, no individuals were found to have ingested plastics (%FO=0%). Similar to the Atlantic salmon sample, the proportion of fed individuals was low at just 37%. Also like Atlantic salmon, capelin are caught while spawning.

### Plastic forensics

The most common plastic type was fragments (*n*=6), followed by threads (*n*=5), then fibres and sheet/film plastics (*n*=4 per category) (Fig. 2A). The particle lengths (longest dimension) ranged from 1.6 mm to 12.1 mm, with a mean length of 5.6 mm (*n*=16, SD=3.140). The majority of particles were microplastics (*n*=11) (Fig. 2B). Size could not be recorded for three plastic particles that were visually identifiable as microplastics, either because they were lost prior to analysis (*n*=2), or were too small to register by the calipers (*n*=1).

**Fig. 2.**
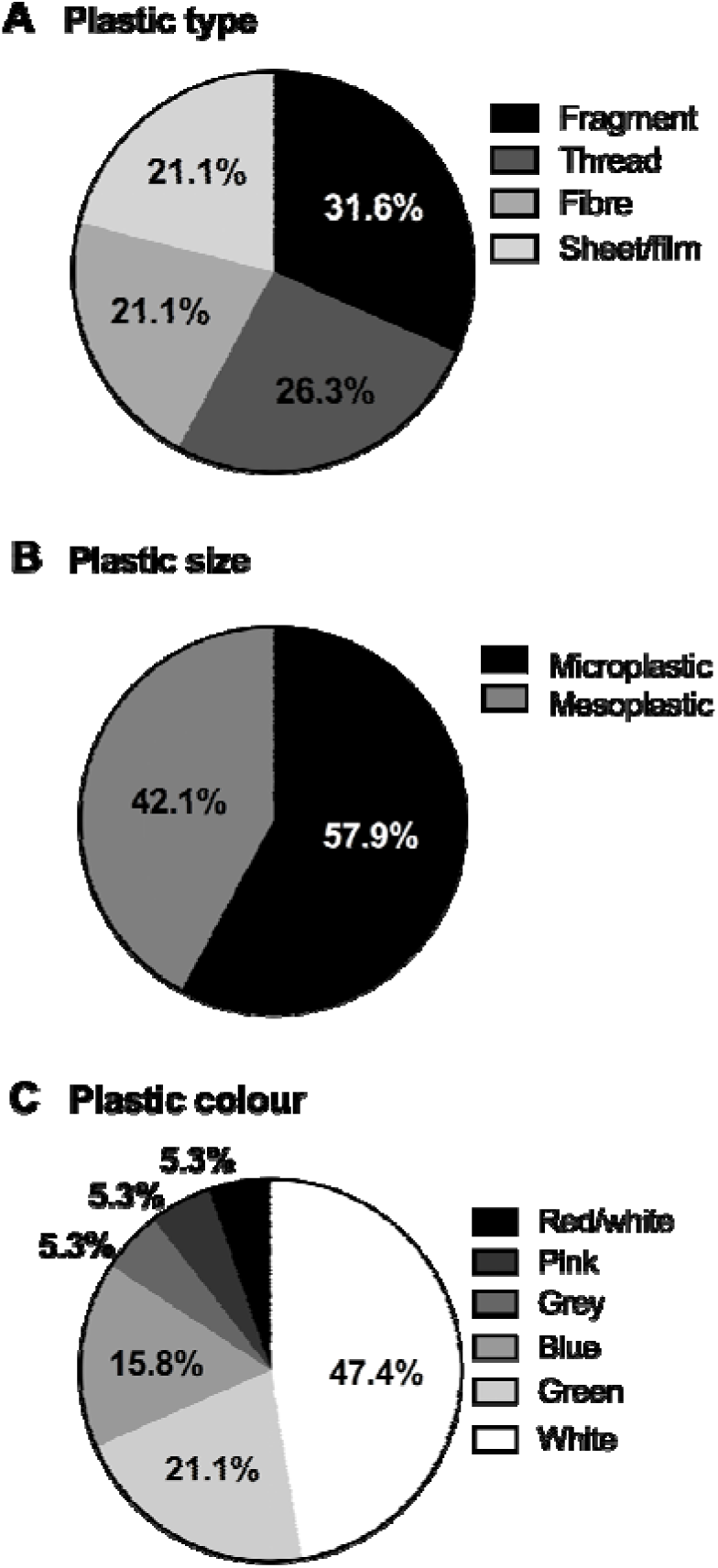
The characteristics for plastic particles (n=19) recovered from Atlantic cod captured in Newfoundland from 2015 to 2016 for the following: (A) plastic type, (B) plastic size, as microplastics (<5 mm) or mesoplastics (5-20 mm) and, (C) plastic colour.

The masses of the plastic for which weight could be determined (*n*=14), ranged from 0.0001 g to 0.6970 g, with a mean mass of 0.0510 g (*n*=14, SD=0.190; Table 4). The masses were not recorded for 5 particles, either because they were lost prior to weighing (*n*=3) or were too lightweight to register (*n*=2). The high standard deviation is attributed to the mass of one of the particles (a large piece of rope); when this particle was removed, the mean mass of the remaining 13 particles was 0.0015 g (*n*=13, SD=0.002). The mean weight of plastic for the entire sample population of Atlantic cod (including individuals that had not ingested plastic) is 0.0007 g per individual (*n*=1,010, SD=0.022). Although this metric is recommended to be reported in ingestion studies for the sake of standardization (Galgani et al., 2013), considering that only 17 cod had ingested any plastic, it is more useful here to report the mean weight of plastics among individuals for which plastic ingestion had occurred. Of the sub-population of Atlantic cod that had ingested plastic(s), the mean weight of plastic is 0.0420 g per individual (*n*=17, SD=0.170).

**Table 4.** Plastic characteristics and measurements for particles recovered from Atlantic cod captured in Newfoundland from 2015 to 2016. Offshore (OS) and inshore (IS) collection sites are noted per location, where offshore was considered >30 km from the shoreline and inshore was <30 km from the shoreline.

| Location | Polymer | Type | Mass (g) | Length (mm) | Colour | Erosion |
| --- | --- | --- | --- | --- | --- | --- |
| Quidi Vidi (IS) | Δ | Thread | 0.0028 | 9.0 | Green | Irregular surface |
| Quidi Vidi (IS) | PVC<sup>a</sup> | Thread | * | 3.5 | White | Fraying |
| St.Philip's (IS) | Δ | Sheet | 0.0017 | 6.5 | White | None |
| St.Philip's (IS) | Δ | Sheet | 0.0011 | 9.7 | White | None |
| St.Philip's (IS) | Δ | Thread | Ø | 8.2 | Green | None |
| St.Philip's (IS) | Ø | Fibre | Ø | Ø | Blue | Fraying |
| Petty Harbour (IS) | Δ | Fragment | 0.0010 | 4.2 | White | Pitting, adhered particles, grooves, linear fractures |
| Petty Harbour (IS) | Δ | Fragment | 0.0002 | 2.0 | White | Pitting, adhered particles, grooves, linear fractures |
| Petty Harbour (IS) | Δ | Fragment | 0.0018 | 2.8 | White | Grooves, linear fractures |
| Witless Bay (IS) | ABS<sup>a</sup> | Fragment | 0.0003 | 1.6 | Grey | None |
| Witless Bay (IS) | PET | Fibre | 0.0015 | 12.1 | Pink | None |
| Bauline East (IS) | PE | Thread | 0.0005 | 6.0 | Green | Small pits, grooves |
| Brigus South (OS) | Δ | Film | 0.0002 | 3.4 | Green | Discoloured, melted, frayed |
| Brigus South (OS) | Ø | Fibre | Ø | Ø | Blue | Fraying, discolouration |
| Halibut Channel (OS) | Δ | Fibre | * | * | Blue | None |
| Burgeo Bank (OS) | PE<sup>a</sup> | Thread | 0.6970 | 9.0 | White | None |
| Grand Banks (OS) | PC<sup>a</sup> | Fragment | 0.0070 | 5.0 | Red/white | None |
| Grand Banks (OS) | PE | Thread | 0.0001 | 4.0 | White | None |
| Grand Banks (OS) | PVC<sup>a</sup> | Fragment | 0.0010 | 3.0 | White | Linear fractures, irregular surface, grooves |

<sup>a</sup> suspected
\* too small to register
Ø particle lost, data could not be gathered
Δ no result

Nine of the particles (47%) had no discernable weathering or erosion patterns, while the rest of particles (53%, *n*=10) had various forms of weathering, including pits and grooves (*n*=5), fracturing (*n*=4), adhered particles (*n*=2), fraying (*n*=4), discolouration (*n*=2), melting (*n*=1) and irregular surface textures (*n*=2) (Table 4). There was a higher proportion of particles recovered from offshore cod that displayed considerable weathering (29%, *n*=2) relative to particles from inshore fish (17%, *n*=2), but the frequency of occurrence of the weathered particles did not vary between offshore and inshore fish (*p*=0.510, df=1, *n*=19).

It is worth noting that three plastic particles were microfibres and the possibility of contamination needs to be addressed. Two particles were enmeshed in a conglomerate of partially digested material (Fig. 3C), eliminating the possibility of contamination via atmospheric deposition, as is common with microfibres (Fries et al., 2013; Hidalgo-Ruz et al., 2012; Nuelle et al., 2014; Woodall et al., 2015). Although the third microfibre was small enough to be atmospherically deposited, it did not match any fibres present in the control dishes and instead was embedded in a bundle of red algae. As such, we do not consider any of these microfibres to be a product of contamination.

**Fig. 3.**
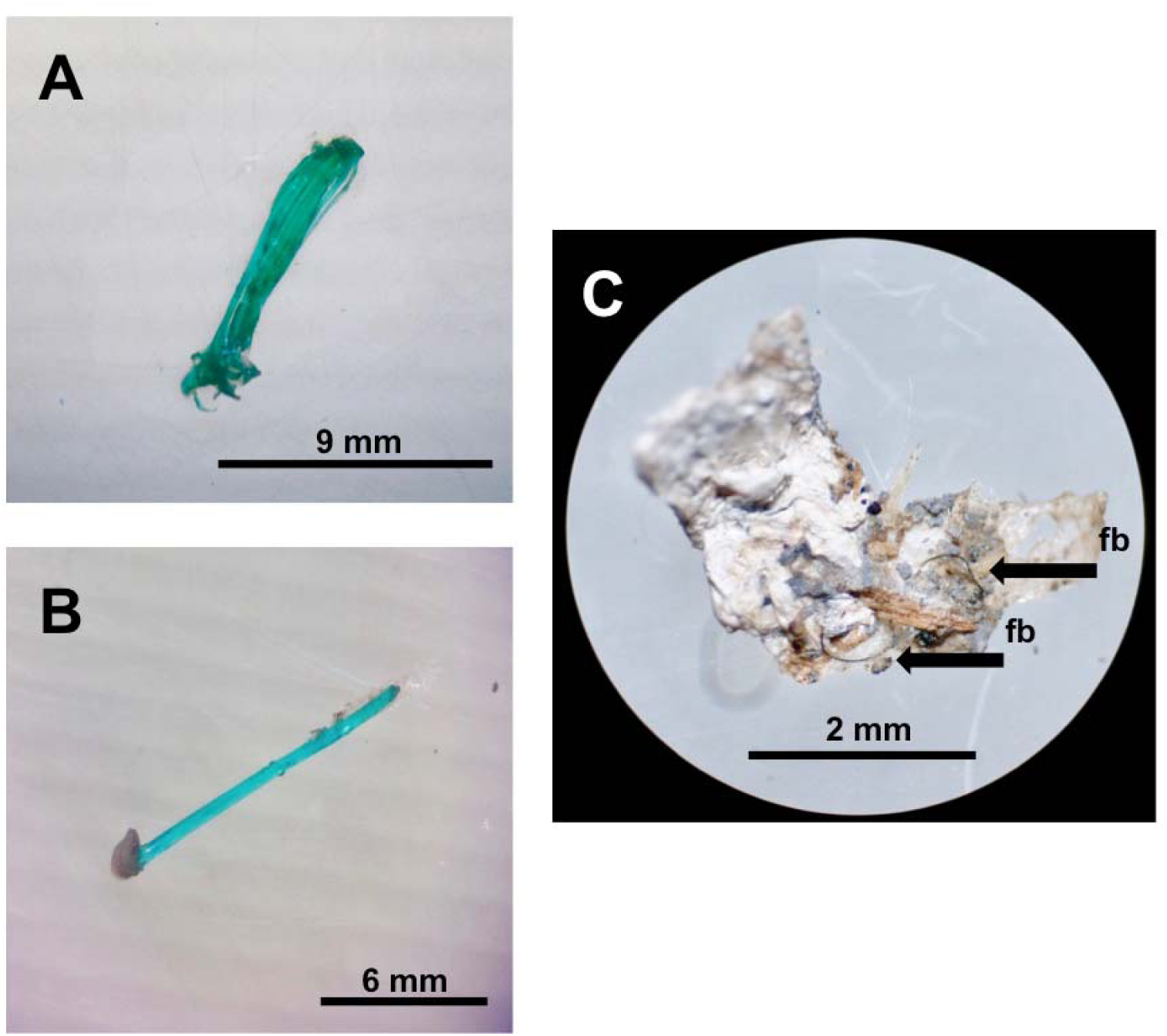
Selection of plastics found in Atlantic cod (*Gadus morhua*) collected by CSS in 2015/2016: (A) green fishing line from 2015, (B) a green polyethylene fishing line thread from 2016 and, (C) a fibre enmeshed in a conglomerate of partially digested material; the black arrows labeled “fb” indicate the exact location of the fibre.

The following plastic polymer types were identified for particles recovered from Atlantic cod using Raman micro-spectrometry: polyethylene (PE), polyvinylchloride (PVC), polyethylene terephthalate (PET), acrylonitrile butadiene styrene (ABS), and polycarbonate (PC). For more details on spectrometry results, see S2.

## Discussion

A total of 1.19% of studied Newfoundland food fish individuals (Atlantic cod, Atlantic salmon and capelin) were found to have ingested plastic, with an ingestion rate of 0% for capelin, 0% for Atlantic salmon, and 1.68% for Atlantic cod. The geographical range of collections was broad, and spanned both the southern and eastern coasts of the island, and northern Labrador (Davis Strait). Atlantic cod is a culturally and economically important species on the island of Newfoundland. In total, 17 individual cod ingested at least one plastic particle.

The majority (58%) of ingested plastics were microplastics (1-5 mm) and the remainder were mesoplastics between 5 mm and 10 mm in size. This is at first a surprising result considering that microplastics are expected to be the most common neuston marine plastic by number (Eriksen et al., 2014), and plastics ingested by fish are often dominated by the microplastic size class (Foekema et al., 2013; Lusher et al., 2016, 2013; Murphy et al., 2017; Neves et al., 2015; Peters et al., 2017). Yet in previous studies, both Bråte et al. (2016) and Rummel et al. (2016) used identical methods to ours to identify plastics in Atlantic cod and in both cases, plastics greater than 5 mm in size were present in the highest proportion.

Half of all ingested particles showed no weathering, and in several particles weathering was minimal. There seemed to be no difference of weathering condition in particles recovered from individuals collected offshore when compared to inshore fish. Particle colours were often fresh and bright, fragments had sharp edges, and most fibres were minimally frayed. This lack of weathering is indicative of short residence times in the ocean, which may point toward a local source (Endo et al., 2005; Liboiron et al., 2016).

Green threads were recovered in Atlantic cod from both sampling years (Fig. 3). This particular shade of green is common in the fishing industry’s most common fishing gear: polyethylene, polypropylene, and polysteel ropes and threads (Knowlton et al., 2016; Murray and Cowie, 2011; Nedostup and Orlov, 2010; Turner, 2016). The green thread recovered from an Atlantic cod individual in 2016 was confirmed by Raman micro-spectrometry to be polyethylene and green was the second most common colour of particles recovered from cod in this research.

All polymers identified in Raman spectrometry (*n*=8) were of a higher density than seawater, except polyethylene (Table 4). A density greater than that of seawater likely leads to the movement of these high density particles from the surface of water to the benthos (Browne et al., 2010; Claessens et al., 2011; Nuelle et al., 2014), placing them in the feeding zone of benthopelagic feeders such as Atlantic cod. Polyethylene was the most commonly identified polymer and is ubiquitously used in the production of fishing gear. Polyethylene in its pristine form is buoyant in seawater (Turner, 2016), though biofouling or ingestion by vertically migrating species can transport this material to greater depths (Lusher et al., 2013; Ye and Andrady, 1991).

Plastics ingestion by Atlantic cod has been examined over a wide geographic region beyond this study, with a particular focus on the Northeast Atlantic. These published results have been highly variable. Rummel et al. (2016) and Bråte et al. (2016) report %FO values of ≤ 3% for Atlantic cod in the North, Baltic and Norwegian seas. By contrast, earlier studies reported higher %FO values for Atlantic cod in the same region; ranging from 13% (Foekema et al., 2013) to 39% (Lenz et al., 2015a) in the North Sea, and 21% in the Baltic Sea (Lenz et al., 2015a). Low plastic ingestion levels reported for Atlantic cod (<3%FO) that are comparable to the results reported here stem from studies that similarly rely on the visual identification of plastics (Bråte et al., 2016; Rummel et al., 2016). Meanwhile, reports of relatively high plastic ingestion levels in Atlantic cod (>10% FO) have stemmed from studies that employ acid digestion of digestive contents and can identify plastics < 1 mm in size (Foekema et al., 2013; Lenz et al., 2015a), thus increasing the size range of identifiable plastics and the resultant %FO for the sample population. Acid digestion protocols are not suitable to the long-term monitoring developed by CLEAR for its inaccessibility to citizen scientists, and so are not used in this study (see Liboiron et al., 2016).

The wide range of sample sizes may also explain the wide range of %FO values reported for Atlantic cod. Outside of this research, sample sizes for Atlantic cod plastic ingestion research range from 7 (Rummel et al., 2016) to 302 (Bråte et al., 2016) individuals. The ideal minimum sample size for plastic ingestion studies is species specific and depends on the %FO for that species (larger sample size required for species exhibiting a lower %FO), as well as any interannual variability in that average and whether the purpose of the research is to detect changes over time (Provencher et al., 2016; van Franeker and Meijboom, 2002). There is not yet enough records of plastic ingestion in Atlantic cod across various sample sizes to conduct an analysis of the species dataset and determine at which sample size the variance in reported %FO decreases, as described by van Franeker and Meijboom (2002).

To our knowledge, this is the first published study that examines plastic ingestion by Atlantic salmon. The majority of salmon examined in our study (84%) were collected within the Newfoundland summer recreational fishing season (Fisheries and Oceans Canada, 2015). During this period, Atlantic salmon return from the ocean to spawn in freshwater rivers in Newfoundland (Porter, 1975). It is possible that individuals were not actively feeding during this time - indeed, only a small proportion (20%) of individuals collected in this study had food in their stomach. A previous study found 1 plastic in 4 individual Chinook salmon destined for human consumption in California, USA, for a %FO of 25%, though the sample size is too low to generalize for the species (Rochman et al., 2015). Ingestion studies on the zooplankton that are food sources for salmon spectated that secondary ingestion of microfibers would be widespread in salmon (Desforges et al., 2015). It is not clear whether these studies are comparable to this one, as the species, methods, and locations are different.

Similar to Atlantic salmon, this is the first study of plastic ingestion by capelin. Sampling of capelin for this study was also restricted to the summer when capelin are subject to human consumption during their spawning period (Lilly, 1987). Similar to Atlantic salmon, the proportion of fed individuals collected in this study was relatively low (37%). The human consumption of capelin may be of particular interest considering that the species is often consumed whole (MacDonald, 2016; Over, 2003).

The 0% ingestion rate in capelin and Atlantic salmon mirror a null rate in Silver hake in the region of southern Newfoundland, as well as 41% of all fish species reported in English-language scientific literature (Liboiron et al., 2018). While the null rate may be due to the species spawning behavior, it may be part of a broader trend.

Geographically, sampling was relatively broad, however, the west and north coasts of Newfoundland as well as coast of Labrador were not examined in this study. Future monitoring of plastics in important food fish species should address these regions.

## Acknowledgements

We acknowledge that this research was conducted on the ancestral Lands of the Mi’kmaq and Beothuk peoples. We would also like to acknowledge the Inuit of Nunatsiavut and NunatuKavut and the Innu of Nitassinan, and their ancestors, as the original peoples of Labrador. We would like to acknowledge Fisheries and Oceans Canada and the Fisheries and Marine Institute of Memorial University for providing the offshore survey samples. We would like to extend our appreciation to the Merschod Lab for help with Raman Spectrometry. We would also like to thank all of the commercial and recreational fishermen who invited us to participate in their filleting process for the donation of samples. For the salmon donations we would like to thank: Dr. Martha Robertson (DFO), Kristin Bøe, Jacqueline Chapman, Robert Lennox and the Cooke Lab. For cod and capelin collections we extend our appreciation to: Centre for Fisheries Ecosystems Research, the crew of the *RV* Celtic Explorer, Susan Fudge, Laura Wheeland and the Government of Newfoundland and Labrador- Fisheries and Land Resources. Finally, we extend special thanks to Coco Coyle and Claudine Metcalf for helping collect and process samples. This this work is based on Masters theses by Jessica Melvin (2017) and Natalie Richárd (forthcoming).

## Supplementary

### S1: Collection methods

The following text outlines collection methods in detail.

#### Scientific surveys

##### Atlantic cod

Atlantic cod were collected in 2016 during the following trawl surveys: 1) Ecosystem Research Program surveys conducted by the Department of Fisheries and Oceans (DFO) aboard the vessel RV *Teleost,* and 2) Transatlantic Added Value survey conducted by the Marine Institute (MI) of Memorial University of Newfoundland aboard the vessel RV *Celtic Explorer.* These surveys are primarily aimed at assessing stock sizes of various species of fish and invertebrates. DFO trawl research evaluates the status of Atlantic cod stocks, while MI uses acoustic and trawl methods to assess different species (Fisheries and Oceans Canada, 2012b; Marine Institute (Foras na Mara), n.d.). Surveys occurred over the southern and eastern Newfoundland shelf, as well as in the Davis Strait.

Both surveys examined individual Atlantic cod for prey analysis prior to our microplastic analyses for an unrelated project. Trained technicians from DFO and MI dissected each stomach from the GI tracts and transferred them to separate, sterile metal trays, where digestive contents were analyzed. Following this analysis, each stomach and all of its contents were placed into a plastic bag with the corresponding GI tract. To ensure all particles were transferred, the metal tray was thoroughly rinsed with a wash bottle and this water was also transferred to the plastic bag, which was then sealed and frozen for later plastic analysis.

##### Atlantic salmon

Atlantic salmon were collected at a fish-counting fence in the Campbellton River. Located 220 m from the mouth of the river, this fence is constructed annually by DFO and used by researchers to study salmon returning to the river (Downton et al., 2001). Atlantic salmon were collected and sacrificed so that physiological samples like blood, isotopes, and fatty acids (not related to the GI tract) could be used for three other research projects. Physiology studies were conducted under Fisheries and Oceans Canada Experimental Licenses provided to Memorial University, Carleton University, and DFO. Donations of intact GI tracts were made for this study after the physiological sampling.

##### Capelin

Capelin were collected by commercial fishers around the island of Newfoundland and donated to this study by DFO. Capelin were caught using several fishing methods (cast net, purse seine, and tuc seine). Whole individuals were frozen immediately after collection and stored for later dissection and analysis.

#### Citizen science surveys

Atlantic cod were collected from the 2015 and 2016 Newfoundland commercial and recreational fisheries. All individuals examined in this survey were collected directly by fishers from six coastal communities located along the Eastern Avalon Peninsula, and the GI tracts were donated to researchers at the wharf when fish were landed. Three of these communities were visited in 2015, and revisited in 2016 (Petty Harbour, Quidi Vidi, and Portugal Cove-St. Philip’s) with the addition of three new communities further south in 2016 (Witless Bay, Bauline East, and Brigus South). Atlantic cod were caught by one of the following methods: hand-line, jigger, or gill net. Such methods are advantageous over scientific trawling as no anthropogenic debris accumulates in the net for possible consumption by captured fish (Davison and Asch, 2011).

In the cod surveys from 2015, around 30 fishers participated per collection site whereas in 2016, between one and seven fishers participated per site. This sampling method is based on fishers’ knowledge of fish aggregations (“good fishing spots”), which increases sampling efficiency

(Liboiron et al., 2016). This sampling method has proved efficient by providing a means of sampling the human food web directly; all fish analysed were consumed by humans (see also Neves et al., 2015; Rochman et al., 2015). The removal of GI tracts (from esophagus to anus) occurred at the wharf and was done either by the researcher or the fisher, depending on the fisher’s preference. Fish were always identified as Atlantic cod by the researcher(s) and other data about individual fish (length, sex) was recorded when possible. All GI tracts were individually bagged and frozen for further analysis.

### S2: Raman Spectrometry

Two of the three microfibres identified in 2016 were lost prior to Raman micro-spectrometry and therefore excluded from the analysis. A total of 10 particles from Atlantic cod caught in 2016, one particle from Atlantic salmon, and two particles from capelin were subjected to Raman micro-spectrometry. Of the 13 particles analysed with Raman micro-spectrometry, three yielded conclusive polymer identifications, five yielded suspected polymer identifications, two particles’ polymers could not be identified (but were visually identifiable as plastics), two were confirmed to be non-plastic and one microfibre was too small to produce a result (Table 4).

**Table S2.** Summary of results for particles subjected to Raman micro-spectrometry analysis. Results indicated whether particles were confirmed as plastic (Y, Yes or N, No), whether a polymer could be identified (Y, Yes or N, No), and, within particles for which a polymer was identified, the confidence of the polymer result (C, conclusive or S, suspect). Dashes indicate information could not be determined in the analysis.

| Species | Particle type | Particle colour | Plastic (Y/N) | Polymer (Y/N) | Confidence (C/S) | Polymer | Density (g/cm<sup>3</sup>)* |
| --- | --- | --- | --- | --- | --- | --- | --- |
| Atlantic cod<br>(*Gadus morhua*) | Fibre | White | Y | Y | S | PVC | 1.15-1.96 |
|  | Fragment | Grey | Y | Y | S | ABS | 0.90-1.53 |
|  | Fibre | Pink | Y | Y | C | PET | 1.38 |
|  | Film | Green | Y | N | - | - | - |
|  | Fibre | Green | Y | Y | C | PE | 0.92-0.96 |
|  | Fibre | Blue | Y | N | - | - | - |
|  | Fibre | White | Y | Y | S | PE | 0.92-0.96 |
|  | Fragment | White+Red | Y | Y | S | PC | 1.36 |
|  | Fibre | White | Y | Y | C | PE | 0.92-0.96 |
|  | Unknown |  | N | - | - | - | - |
|  | Unknown |  | N | - | - | - | - |
|  | Fragment | White | Y | Y | S | PVC | 1.15-1.96 |
| Capelin<br>(*Mallotus villosus*) | - | - | N | - | - | - | - |
|  | - | - | N | - | - | - | - |
| Atlantic salmon<br>(*Salmo salar*) | - | - | N | - | - | - | - |

\* Densities listed here correspond to the pure polymer, according to Wolfram Alpha LLC (2017). Actual particle densities will vary from that of their pure form. Seawater density ~ 1.02-1.04 g/cm<sup>3</sup> (Wolfram Alpha LLC, 2017)

